# Crebinostat Facilitates Memory Formation

**DOI:** 10.1101/2024.02.05.578875

**Authors:** Deepti Dama, Shiv K Sharma

## Abstract

**Objectives:** Protein modifications importantly contribute to memory formation. Protein acetylation is a posttranslational modification of proteins that regulates memory formation. Acetylation level is determined by the relative activities of acetylases and deacetylases. Crebinostat is a histone deacetylase inhibitor. In this study, our aim was to examine whether Crebinostat affects memory formation in object recognition task. Further, we aimed to understand regulation of plasticity-related proteins by Crebinostat.

**Methods:** Object recognition task was used to examine the effect of Crebinostat on memory formation. The effect of Crebinostat on plasticity-related proteins was examined using Western blot experiments.

**Results:** We show that Crebinostat facilitates memory formation by a weak training in object recognition task. Further, this compound enhances acetylation of α-tubulin, and reduces the level of histone deacetylase 6, an α-tubulin deacetylase.

**Conclusions:** The results suggest that enhanced acetylation of α-tubulin by Crebinostat contributes to its facilitatory effect on memory formation.

**Abstract:** Protein modifications importantly contribute to memory formation. Protein acetylation is a posttranslational modification of proteins that regulates memory formation. Acetylation level is determined by the relative activities of acetylases and deacetylases. Crebinostat is a histone deacetylase inhibitor. Here we show that in object recognition task, Crebinostat facilitates memory formation by a weak training. Further, this compound enhances acetylation of α-tubulin, and reduces the level of histone deacetylase 6, an α-tubulin deacetylase. The results suggest that enhanced acetylation of α-tubulin by Crebinostat contributes to its facilitatory effect on memory formation.

## Introduction

The post-translational modifications in proteins are known to play critical roles in synaptic plasticity and memory formation [1]. For example, phosphorylation that is regulated by kinases and phosphatases, regulates these processes. The observation that long-term memory (LTM) requires new gene expression, led to the identification of transcription factors that regulate expression of memory-related genes. Since the status of chromatin also regulates transcriptional processes [2], this aspect also has received significant attention for its role in memory formation. Histones undergo several post-translational modifications including acetylation [3, 4]. Acetylation is a reversible modification in which histone acetylases (HATs) and histone deacetylases (HDACs) add or remove the acetyl group, respectively. Histone acetylation in the chromatin is typically associated with transcriptional activation [5].

It has become clear that acetylation critically regulates synaptic plasticity and memory [6, 7]. For example, Levenson and colleagues showed that histone acetylation is enhanced 4 by memory training [8]. Since the level of acetylation is regulated by the relative activities of acetylases and deacetylases, inhibition of deacetylases has served as a useful tool to increase acetylation levels. Several studies have used this approach to study the effect of increasing acetylation levels on memory formation. Levenson and colleagues showed that increasing acetylation levels by inhibition of histone deacetylases enhances memory [8]. Likewise, several other studies have shown a positive effect of increasing acetylation on synaptic plasticity and memory [9, 10, 11, 12]. Importantly, increasing acetylation levels helps recovery of memory in a model of neurodegeneration [13]. Further, increasing acetylation levels has shown beneficial effects in Alzheimer’s disease, and in age-related memory decline [14, 15].

Crebinostat was identified as a histone deacetylase inhibitor and an activator of cAMP response element-binding protein (CREB)-mediated transcription [16]. This compound enhances memory formation in a contextual fear memory task [16]. Here we have examined the effect of Crebinostat on memory formation by a weak training in novel object recognition paradigm. Further, we have examined its effects on plasticity-related proteins.

## Materials and Methods

### Animals

Sprague-Dawley male rats were used for the experiments. For behavioural experiments, 10–12-week-old animals were used, and for molecular studies, 6–8-week-old animals were used. The animals were housed in the National Brain Research Centre’s animal facility on a 12-hour light / 12-hour dark cycle with ad libitum access to animal feed and water. The procedures were approved by the Institutional Animal Ethics Committee of National Brain Research Centre.

### Behavioural experiments

The experiments used novel object recognition task to examine memory. This task was performed between 6.30 pm and 10.30 pm. The animals were habituated to the arena for 4 days during which they were allowed to freely explore the arena devoid of any objects for a 10 min duration, each day. During training, the animals were exposed to two replicas of an object, A1 and A2. Twenty-four after the training, long-term memory for the object was tested. During the LTM test, the animals were allowed 5 min to explore a new object, B, and one of the objects that was used during training. For the 10-min training group, during training, the animals were given 10 min to explore the objects. The Crebinostat group of animals were given intraperitoneal injection of Crebinostat (25 mg/kg in 10% DMSO, 20% Cremophore, and 70% saline) 1 h before training during which they were exposed to the objects for 5 min. The animals in the control group received vehicle (10% DMSO, 20% Cremophore and 70% saline) 1 h before training that consisted exposure to the objects for 5 min. The interaction with each object was scored if the snout of the animal was facing towards the object while it was smelling, licking, or biting it for a duration of 2 sec or longer. ANYMAZE software was used to record the animal activities, and the recorded videos were used to determine the time of interaction manually. A discrimination index (DI) for each animal was determined using the formula, DI = interaction time with the new object / total time of interaction with both the objects [17].

### Hippocampal slicing and treatment

The animals were anaesthetized using Halothane (Piramal) and killed. The brains were dissected out. Hippocampi were isolated, and transverse sections of 350-400 µm thickness were obtained using the protocol described previously [18]. After recovery, the slices were randomly divided into two groups with 3-4 slices in each group. One group of slices was treated with Crebinostat [25 µM Crebinostat in artificial cerebrospinal fluid (aCSF: 125 mM NaCl, 2.5 mM KCl, 1.25 mM NaH_2_PO_4_, 25 mM NaHCO_3_, 2 mM CaCl_2_,1 mM MgCl_2_, 25 mM glucose)] for 1 h. The stock solution of Crebinostat was prepared at a concentration of 25 mM in DMSO. The control group of slices was treated with DMSO in aCSF for the same duration. Post-treatment, the slices were rinsed with aCSF and collected in sodium dodecyl sulphate (SDS) lysis buffer (50 mM Tris-Cl, pH 7.5, 150 mM NaCl, 50 mM NaF, 1 mM sodium orthovanadate, 10 mM sodium butyrate, 1 mM EDTA, 2% SDS, protease inhibitor cocktail).

### Western blotting

Protein extract was prepared from slices and protein estimation was done using the BCA method (ThermoFisher Scientific BCA Kit). Samples with equal amounts of protein were resolved on a polyacrylamide gel. The proteins were transferred to a nitrocellulose membrane which was then blocked with 5% bovine serum albumin in Tris-buffered saline (TBST: 25 mM Tris-Cl, 137 mM NaCl, 2.7 mM KCl and 0.1% Tween-20; pH was adjusted to 7.5 before adding Tween-20) and incubated with primary antibodies [α-tubulin: catalogue no. 2125S (Cell Signalling Technology), acetylated α-tubulin: catalogue no. SC-23950 (Santa Cruz Biotechnology), HDAC6: catalogue no. D21B10 (Cell Signalling Technology), β-tubulin: catalogue no. 2146 (Cell Signalling Technology)] overnight at 4° C. The blots were washed with TBST and probed with HRP-conjugated secondary antibody and incubated for 1 h at room temperature. The blots were again washed with TBST. Signals were detected using BioRad Gel Doc XR imaging system or Uvitec Gel Documentation System. The HDAC6 antibody recognized a band at approximately 130 kDa. For acetylation of α-tubulin analysis, the blots were probed with acetyl α-tubulin antibody, stripped and re-probed with α-tubulin antibody. For the analysis of HDAC6 level, since there is a considerable difference between the molecular weights of HDAC6 and β-tubulin, the blots were cut and probed with the respective antibodies. Bands were quantified using Quantity One or Uvitec-1D software. For estimation of α-tubulin acetylation, the acetyl-α-tubulin signal was divided by the α-tubulin signal in each sample, then normalized with the control sample. Similarly, for estimation of HDAC6 level, the HDAC6 signal was divided by the β-tubulin signal and then normalized with the control sample.

### Data Analysis

During training, for each group, a two-tailed paired Student’s t-test was used to assess the statistical significance of the difference in time of exploration of the two objects. A One-Sample t-test was used to assess the statistical significance of the discrimination index, for each group. An unpaired t-test was used on the discrimination index to assess the effect of Crebinostat. A two-tailed paired t-test was used on acetyl-α-tubulin/α-tubulin signal ratios or on HDAC6/β-tubulin signal ratios to assess the statistical significance of the difference in the two groups. A p-value of less than 0.05 was considered statistically significant. Data are shown as mean +/-SEM.

## Results

### Crebinostat facilitates long-term memory after a weak training

Novel object recognition task is a simple and robust task to assess memory. A 10 min training in this task leads to LTM formation [17, 19]. We first reproduced this finding. After habituation, the animals were exposed to two copies (A1 and A2) of an object for a period of 10 min. During training, there was no significant difference in the exploration time of the objects (Fig. 1A). Twenty-four hours after the training, a probe trial was conducted to assess LTM. During the probe trial, the animals explored a new object, B, and one object that was present during training. The animals spent more time exploring the new object compared to the old object used during training as indicated by the discrimination index (Fig. 1B).

**Figure 1:**
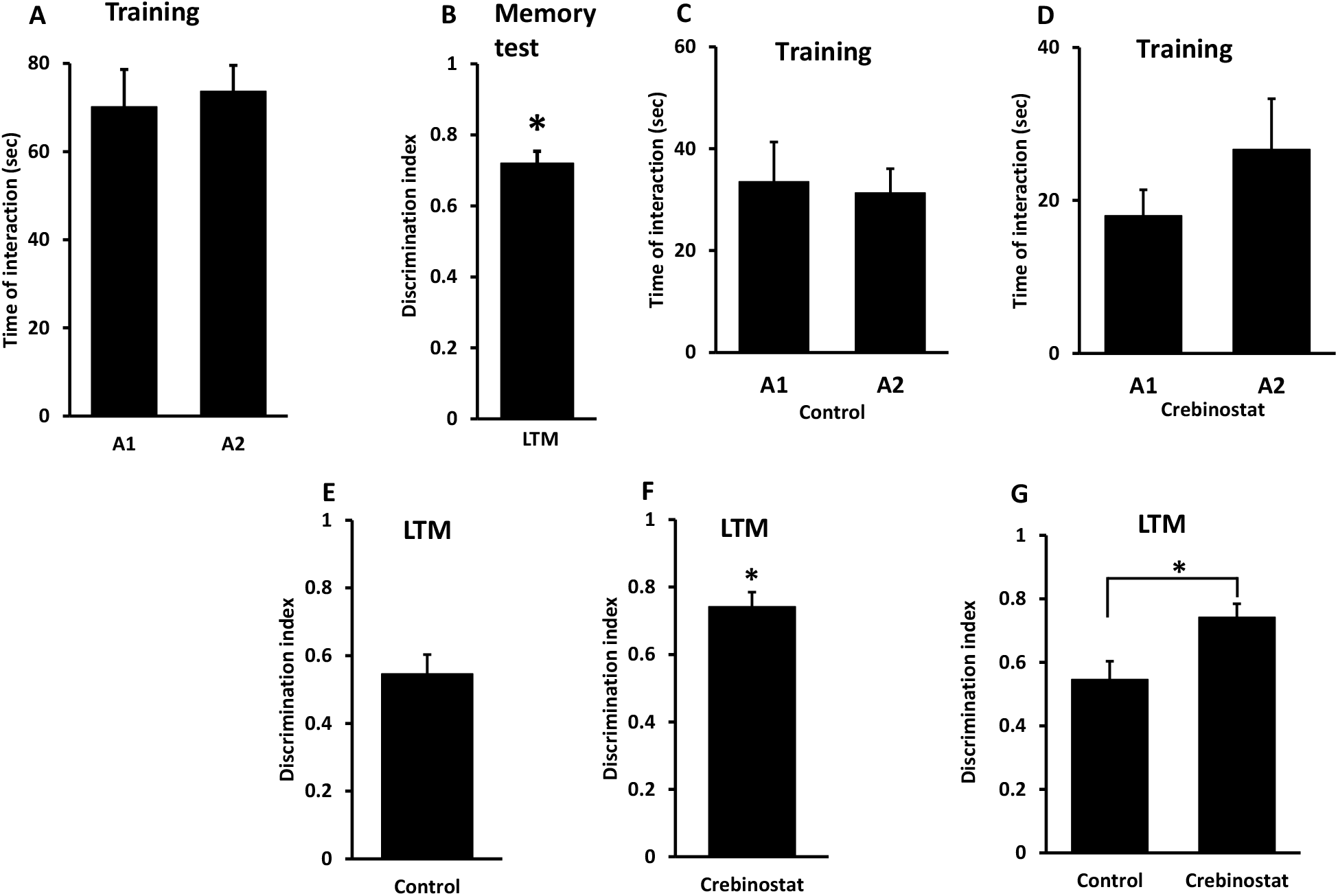
Long-term memory formation in object recognition task (A - B) and crebinostat facilitates memory formation in object recognition task (C - G). For the 10-min training group, during training, the animals were exposed to two copies (A1 and A2) of an object for 10 min. LTM was tested 24 h after the training. During training, the animals spent similar amount of time exploring the two objects (A; n = 6). Discrimination index shows that during the LTM test, these animals preferred the new object compared to the old object used during training (B). To examine the effect of crebinostat on memory formation, the animals were given intraperitoneal injection of either vehicle (Control) or crebinostat (Crebinostat) before training. During training, the animals were exposed to two copies (A1 and A2) of an object for 5 min. LTM was tested 24 h after the training. During training, the control group (C) or the crebinostat group (D) spent similar amount of time exploring A1 and A2 objects (n = 6 in both groups). As indicated by the discrimination index, during the LTM test, the control group did not show any preference for the new object (E), however, the crebinostat group spent more time exploring the new object than the old object used during training (F). A significant difference was observed in the discrimination index between the control and the crebinostat groups (G). A discrimination index of 0.5 indicates that the animals did not prefer the new object over the old object. Asterisks denote a significant difference (p < 0.05).

**Figure 2:**
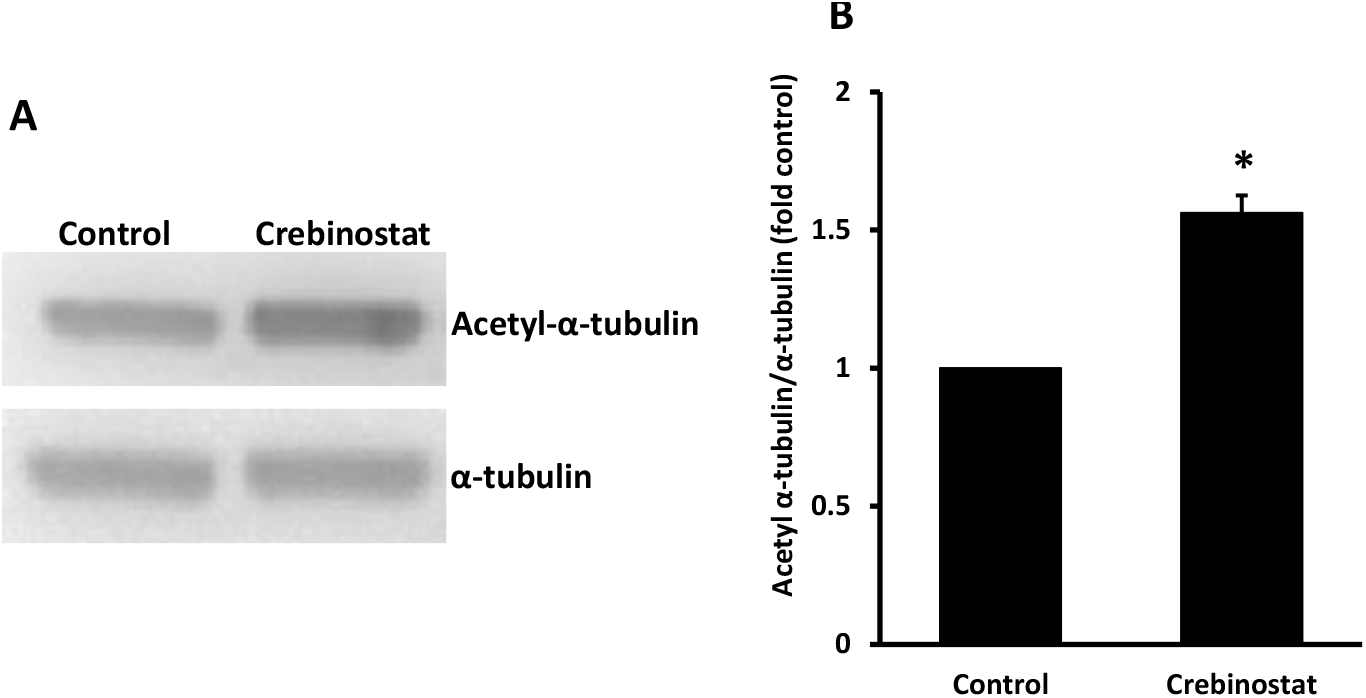
Crebinostat enhances α-tubulin acetylation. The hippocampal slices were treated with Crebinostat (Crebinostat) or vehicle (Control). The blots were probed with acetyl-α-tubulin and α-tubulin antibodies. The representative blots (A) and quantified summary of data (B, n = 5) show that Crebinostat significantly increased acetylation of α-tubulin in the hippocampal slices. Asterisk denotes significant difference (p < 0.05) from control.

We next asked whether Crebinostat has any effect on memory formation in this task. The animals received intraperitoneal injection of either vehicle or crebinostat 1 h before training. A previous study has shown that intraperitoneally-injected crebinostat reaches the brain [16]. During the training phase, they were exposed to two copies (A1 and A2) of an object for 5 min. During training, there was no significant difference in the time of exploration of the two objects by the two groups (Fig. 1C and 1D). Although the Crebinostat group explored object A2 more than object A1, this difference was not statistically significant. Twenty-four hours after the training, a probe trial for LTM was conducted. The discrimination index shows that, during the LTM test, the control group did not show a preference for the new object (Fig. 1E) whereas the Crebinostat group preferred the new object over the old object used during training (Fig. 1F). In addition, there was a significant difference in the discrimination index between the Crebinostat and the control groups (Fig. 1G, p < 0.05). Further, the discrimination index in the Crebinostat group was similar to the discrimination index for the group of animals that had received 10-min training in a separate experiment (Fig. 1B). Thus, although a weak 5-min training does not induce LTM in the object recognition task, Crebinostat facilitates LTM formation with this weak training.

### Crebinostat increases acetylation of α-tubulin

In a previous study, we showed that α-tubulin acetylation is enhanced by memory training [17]. Thus, we asked whether Crebinostat affects α-tubulin acetylation. The slices were given Crebinostat or vehicle treatment. The samples were processed using acetyl-α-tubulin and α-tubulin antibodies. The results show that Crebinostat significantly enhanced α-tubulin acetylation (Fig. 3). This is in agreement with the findings of Fass and colleagues [16] who showed that in an in vitro reaction, Crebinostat inhibits HDAC6, a tubulin deacetylase.

**Figure 3:**
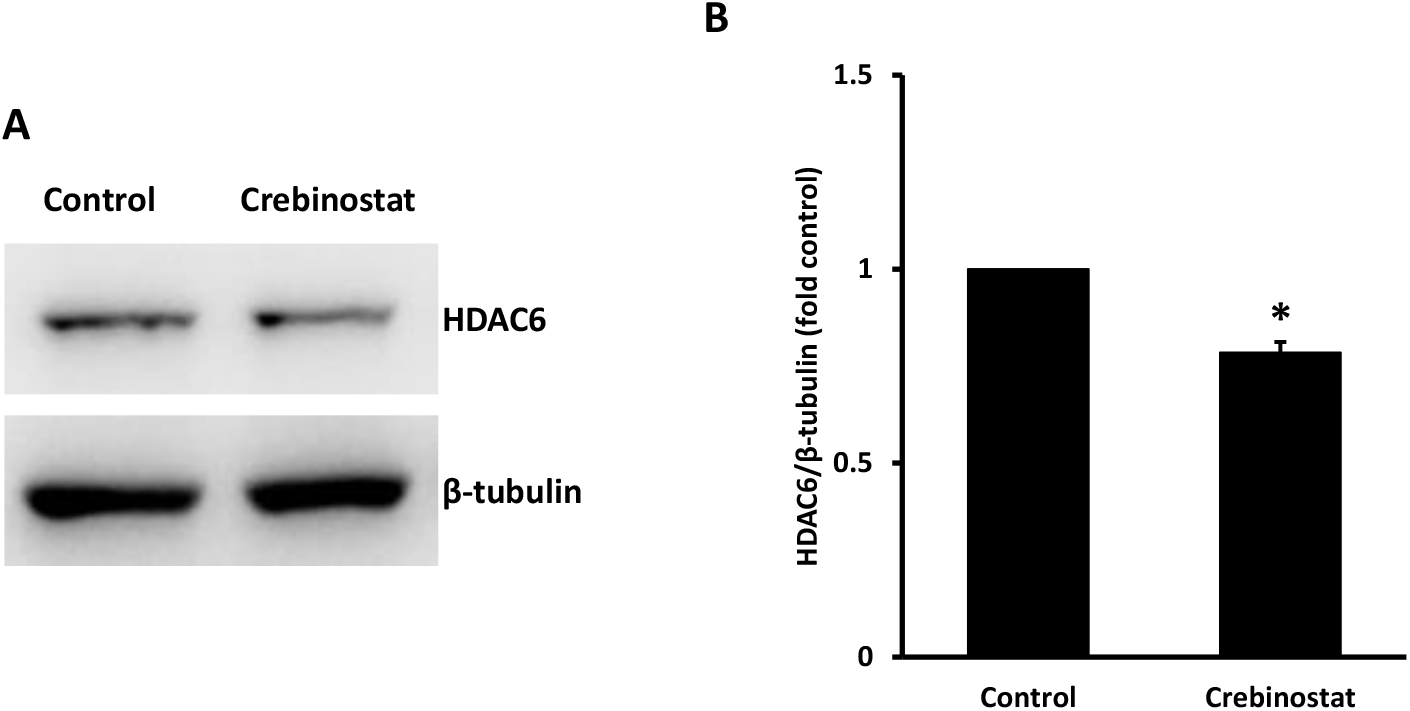
Crebinostat reduces HDAC6 level. The hippocampal slices were treated with crebinostat (Crebinostat) or vehicle (control). The blots were probed with HDAC6 and β-tubulin antibodies. Representative blots (A) and quantified summary of data (B, n = 5) show that Crebinostat significantly reduced HDAC6 level in the hippocampal slices Asterisk denotes a significant difference (p < 0.05) from control.

### Crebinostat reduces HDAC6 level

The results described in the previous section establish that Crebinostat enhances α-tubulin acetylation. Considering that HDAC6 is a tubulin deacetylase, we asked whether HDAC6 level is affected by Crebinostat. The slices were given Crebinostat or vehicle treatment, and the samples were processed using HDAC6 antibody and β-tubulin antibody. The results show that Crebinostat significantly reduced the level of HDAC6 (Fig. 4). Thus, the effect of Crebinostat on α-tubulin acetylation may involve HDAC6 inhibition and a reduction in its level.

## Discussion

Long-term memory formation critically requires new transcription and protein synthesis [20, 21]. In addition to transcription factors, chromatin modifications also play an important role in transcriptional process [5, 22]. Histone acetylation is a well-studied chromatin modification that regulates transcription [23]. Studies have shown that increasing acetylation levels enhances memory [7, 24]. A common approach to increase acetylation is to inhibit the deacetylases. Crebinostat is an HDAC inhibitor that activates CREB-mediated transcription [16]. Further, this compound enhances memory formation in contextual fear memory task [16].

In this study, we investigated the effect of Crebinostat on memory formation using object recognition task. The results show that this compound facilitates LTM by a training paradigm that normally does not induce this form of memory. These results are broadly consistent with the results of Fass and colleagues [16] who showed that administration of Crebinostat for over a week enhances memory in a contextual fear conditioning task. Our results suggest that administration of a single dose of Crebinostat 1 h before training is sufficient to induce LTM by a weak training.

We further examined the regulation of plasticity-related proteins by Crebinostat. Our previous study has shown that acetylation of α-tubulin is regulated by neuronal depolarization [25]. In another study, we showed that acetylation of α-tubulin is increased by memory training in object recognition task [17]. In this study, we found that Crebinostat enhances acetylation of α-tubulin in the hippocampus. Fass and colleagues [16] have shown that Crebinostat inhibits HDAC6, a tubulin deacetylase, in in-vitro conditions. The results show that this compound enhances α-tubulin acetylation in the hippocampal slices. A relatively short treatment of 1 h was sufficient to enhance α-tubulin acetylation in the hippocampal slices.

Since HDAC6 is a deacetylase that regulates α-tubulin acetylation, it is expected that its inhibition by Crebinostat would increase α-tubulin acetylation. We find that HDAC6 level itself is regulated by Crebinostat in the hippocampal slices. It is unclear how crebinostat reduces the level of HDAC6. Further experiments are needed to examine the mechanisms of crebinostat-mediated reduction in HDAC6 level. Inhibition of HDAC6 and a reduction in its level by Crebinostat could contribute to increased acetylation of α-tubulin by Crebinostat. Considering that α-tubulin acetylation is regulated by memory training [17], the findings of this study suggest that one way by which Crebinostat enhances memory formation after a weak training in object recognition task is by increasing the level of α-tubulin acetylation. Inhibition of other HDACs by crebinostat could also contribute to its effect on facilitation of memory. Further studies are needed to fully elucidate the mechanisms by which Crebinostat facilitates memory formation. In addition, it would be interesting to examine the effects of Crebinostat on long-term potentiation which is considered to be a cellular mechanism for memory formation.

## Conclusions

This study shows that Crebinostat facilitates LTM in object recognition task by a weak training which by itself does not lead to LTM formation. This compound increases acetylation of α-tubulin, and reduces the level of HDAC6, an α-tubulin deacetylase. It would be interesting to elucidate the mechanisms involved in facilitation of memory formation by Crebinostat.

## Author contributions

DD and SKS designed the study. DD performed the experiments and analysed the data. DD and SKS wrote the manuscript.

## Acknowledgements

This study was supported by a core grant from Department of Biotechnology, India to National Brain Research Centre. We thank Biswaranjan Sahoo and Apoorva Agrawal for their valuable input.

## Notes

**Conflict of interest statement:** The authors declare no conflict of interest.

**Funding:** This work was supported by a core grant from Department of Biotechnology, India to National Brain Research Centre.

### Competing Interest Statement

The authors have declared no competing interest.

